# A Principal Odor Map Unifies Diverse Tasks in Human Olfactory Perception

**DOI:** 10.1101/2022.09.01.504602

**Authors:** Brian K. Lee, Emily J. Mayhew, Benjamin Sanchez-Lengeling, Jennifer N. Wei, Wesley W. Qian, Kelsie Little, Matthew Andres, Britney B. Nguyen, Theresa Moloy, Jane K. Parker, Richard C. Gerkin, Joel D. Mainland, Alexander B. Wiltschko

## Abstract

Mapping molecular structure to odor perception is a key challenge in olfaction. Here, we use graph neural networks (GNN) to generate a Principal Odor Map (POM) that preserves perceptual relationships and enables odor quality prediction for novel odorants. The model is as reliable as a human in describing odor quality: on a prospective validation set of 400 novel odorants, the model-generated odor profile more closely matched the trained panel mean (n=15) than did the median panelist. Applying simple, interpretable, theoretically-rooted transformations, the POM outperformed chemoinformatic models on several other odor prediction tasks, indicating that the POM successfully encoded a generalized map of structure-odor relationships. This approach broadly enables odor prediction and paves the way toward digitizing odors.

**One-Sentence Summary:** An odor map achieves human-level odor description performance and generalizes to diverse odor-prediction tasks.

## Introduction

A fundamental problem in neuroscience is mapping the physical properties of a stimulus to perceptual characteristics. In vision, wavelength maps to color; in audition, frequency maps to pitch. By contrast, the mapping from chemical structures to olfactory percepts is poorly understood. Detailed and modality-specific maps like the CIE color space (*1*), and Fourier space (*2*) led to a better understanding of visual and auditory coding. Similarly, to better understand olfactory coding, olfaction needs a better map.

Pitch increases monotonically with frequency; in contrast, the relationship between odor percept and odorant structure is riddled with discontinuities, exemplified by Sell’s triplets (*3*), trios of molecules in which the structurally similar pair is not the perceptually similar pair (Fig. 1A). These discontinuities in the structure-odor relationship suggest that standard chemoinformatic representations of molecules—functional group counts, physical properties, molecular fingerprints, etc.— used in recent odor modeling work (*4*–*6*) are inadequate to map odor space.

**Fig. 1.**
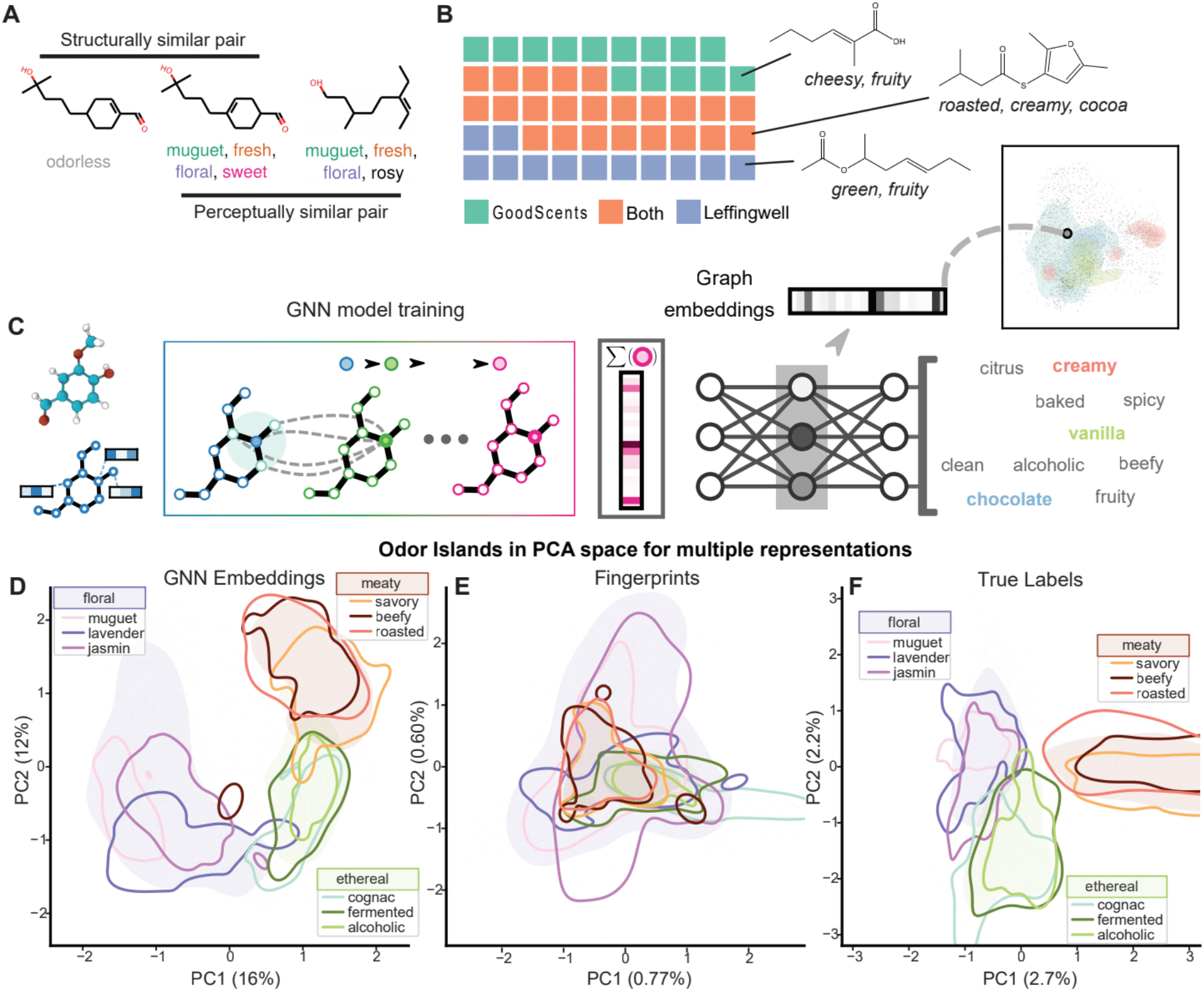
POM preserves the structure of odor perceptual space. **(A)** Example triplet of molecules in which the structurally similar pair is not the perceptually similar pair. **(B)** The GNN was trained on a curated dataset of ~5000 semantically labeled molecules drawn from GoodScents (*13*) and Leffingwell (*14*) flavor and fragrance databases; one square represents 100 molecules; three example training set molecules and their odor descriptions are shown: 2-methyl-2-hexenoic acid (top), 2,5-dimethyl-3-thioisovalerylfuran (middle), 1-methyl-3-hexenyl acetate (bottom). **(C)** Schematic illustrating the process of training a GNN to generate the POM. **(D-F)** Odorants plotted by the first and second principal components (PC) of their **(D)** perceptual labels from GS/LF training dataset (138 labels), **(E)** cFP structural fingerprints (radius 4, 2048-bit), and **(F)** POM coordinates (256 dimensions). Areas dense with molecules having the broad category labels floral, meaty, or alcoholic are shaded; areas dense with narrow category labels are outlined. The POM recapitulates the true perceptual map, but the FP map does not; note that only relative (not absolute) coordinates matter.

## Results

To generate odor-relevant representations of molecules, we constructed a Message Passing Neural Network (MPNN) (*7*), a specific type of graph neural network (GNN) (*8*), to map chemical structures to odor percepts. Each molecule is represented as a graph, with each atom described by its valence, degree, hydrogen count, hybridization, formal charge, and atomic number. Each bond is described by its degree, aromaticity, and whether it is in a ring. Unlike traditional fingerprinting techniques (*9*), which assign equal weight to all molecular fragments within a set bond radius, a GNN can optimize fragment weights for odor-specific applications. Neural networks have unlocked predictive modeling breakthroughs in diverse perceptual domains (e.g., natural images (*10*), faces (*11*), and sounds (*12*)) and naturally produce intermediate representations of their input data that are functionally high-dimensional, data-driven maps. We use the final layer of the GNN (henceforth, “our model”) to directly predict odor qualities, and the penultimate layer of the model as a principal odor map (POM). The POM 1) faithfully represents known perceptual hierarchies and distances, 2) extends to novel odorants, 3) is robust to discontinuities in structure-odor distances, and 4) generalizes to other olfactory tasks.

To train the model, we curated a reference dataset of approximately 5000 molecules, each described by multiple odor labels (e.g. creamy, grassy), by combining the Goodscents (*13*) and Leffíngwell (*14*) (GS/LF) flavor and fragrance databases (Fig. 1B). The model (Fig. 1C) achieved strong cross-validation predictive performance of AUROC=0.89 (*15*).

To test how well the POM represents known perceptual relationships, we compared both the POM and a map built with standard chemoinformatic features - Morgan fingerprints (FP) - to empirical perceptual space (Fig. 1D-F). We measured the fidelity of the maps in representing true relative perceptual distances, (e.g. two molecules that smell of jasmine should be nearer to each other than to a beefy molecule) and hierarchies (e.g. jasmine and lavender are subtypes of the floral odor family). The POM better represents relative distances: distances in the perceptual map (Fig. 1D) are more significantly correlated to distances in the POM (R=0.73, Fig. S1A) than to distances in the FP map (R=-0.12, p <0.001, Fig. S1B). The POM better represents perceptual hierarchies: molecules with a shared odor label have significantly tighter cluster density (CD) in the POM (CD = 0.51± 0.19) than in the FP map (CD = 0.68 ± 0.23, p <0.001, Fig. S2), where smaller CD values denote more dense clusters.

To test if the model extends to novel odorants, we designed a prospective validation challenge (*16*) in which we benchmarked model predictive performance against individual human raters. In olfaction, no reliable instrumental method of measuring odor perception exists, and trained human sensory panels are the gold standard for odor characterization (*17*). Like other sensory modalities, odor perception is variable across individuals (*18*, *19*), but group-averaged odor ratings have been shown to be stable across repeated measurements (*20*) and represent our best avenue to establish the ground-truth odor character for novel odorants. We trained a cohort of subjects to describe their perception of odorants using the Rate-All-That-Apply method (RATA) and a 55-word odor lexicon. During training sessions, each term in the lexicon was paired with visual and odor references (Table S1; Fig. S3). Only subjects that met performance standards on the pretest of 20 common odorants (Data S2; individual test-retest correlation R > 0.35; reasonable label selection for common odorants) were invited to join the panel.

To avoid trivial test cases, we applied the following selection criteria for the set of 400 novel odorants: 1) molecules must be structurally distinct from each other (Fig. S4), 2) molecules should cover the widest gamut of odor labels (Data S1), and 3) molecules must be structurally or perceptually distinct from any training example (e.g. Fig. 1A, Data S1). Our prospective validation set consists of 55-odor label RATA data for 400 novel, intensity-balanced odorants generated by our cohort of ≥15 panelists (2 replicates). Summary statistics and correlation structure of the human perceptual data is presented in Fig. S5-7. Our panel’s mean ratings were highly stable (panel test-retest: R = 0.80, n = 15; Fig. S8) and more consistent than the DREAM cohort’s ratings (*6*) (Fig. S9-10).

Of the 400 molecules characterized, 80 were dropped from the final prospective validation set due to low intensity (42) (Fig. S11), redundancy (1), mistaken inclusion (1), or with confirmed or potential contamination (26) (Data S1). Model performance was evaluated on the remaining 320 molecules without model retraining.

To measure the model’s performance, we compared the concordance of its normalized predictions with the normalized panel mean rating (Fig. 2A and 2C). While there is considerable variation across molecules in the ability of both individual raters and the model to match the panel mean ratings, the model output comes closer to the panel mean than does the median panelist for 53% of molecules (Fig. 2E and 2F). The model’s superiority at the task is even more impressive given that panelists are able to smell each odorant as they rate it, while the model’s predictions are based solely on nominal molecular structure.

**Fig. 2:**
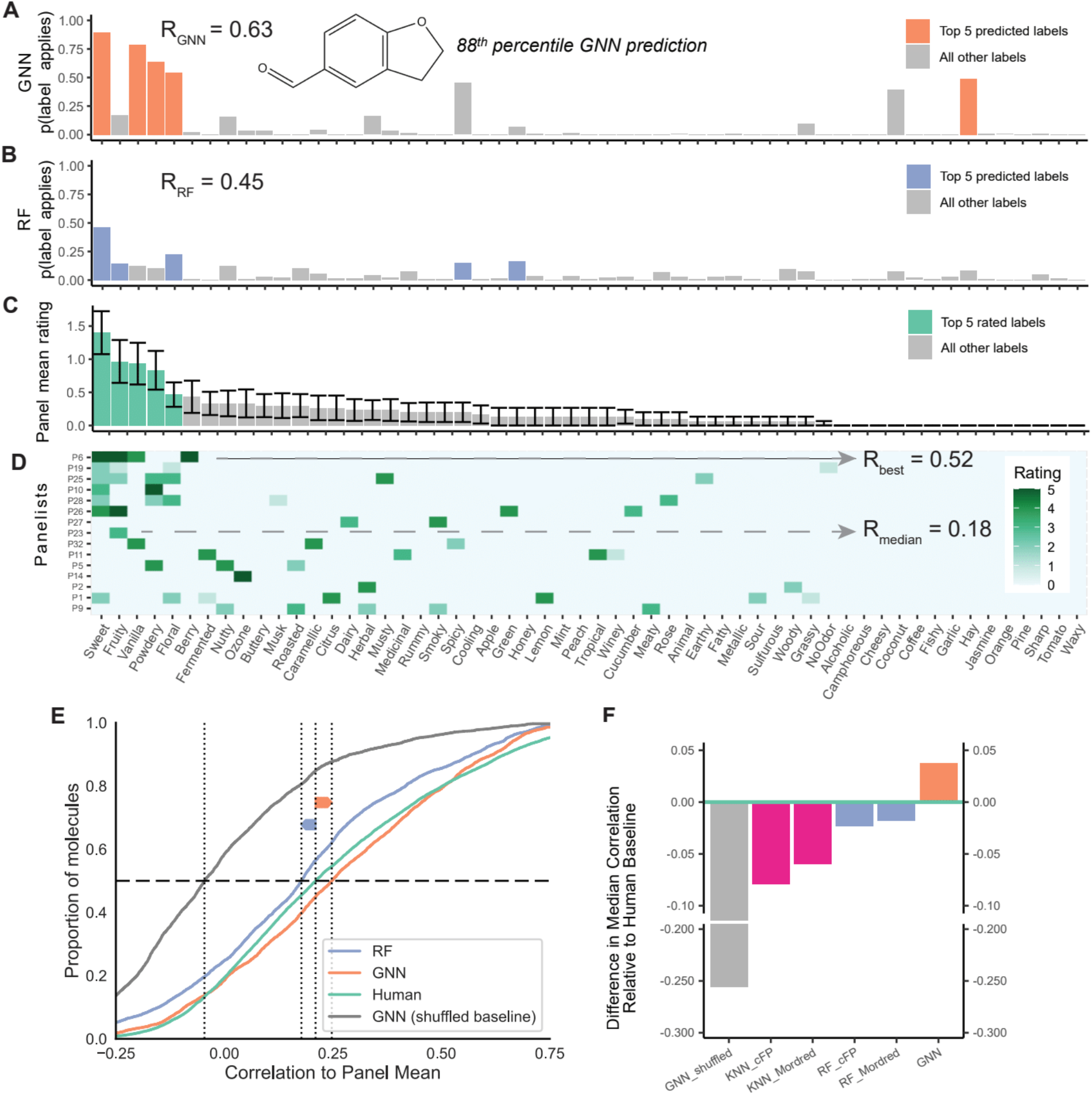
GNN model displays human-level odor description performance. **(A)** GNN model label predictions, **(B)** random forest (RF) model label predictions, **(C)** panel mean ratings with standard error bars, and **(D)** individual panelist ratings, averaged over 2 replicates, for the molecule 2,3-dihydrobenzofuran-5-carboxaldehyde. In panels **A-C**, the top 5 ranked descriptors are in orange (GNN), purple (RF), or green (panel). Descriptors in panels **A-D** are ordered by panel mean ratings. Panels **A**, **B**, and **D** are annotated with the Pearson correlation coefficient of their data to the panel mean rating shown in panel **C**. Panel **D** includes panelist/panel correlation coefficients for the panelist that best matches the panel mean and for the panelist with the median match. **(E)** Cumulative density plot showing the distribution of correlations between human panelists and the panel mean (in green) and between the GNN, RF, and GNN shuffled model predictions and the panel mean on a per molecule basis. Curves shifted to the right are more strongly correlated to the panel mean. **(F)** Difference in the median correlation to the panel mean relative to the median human subject’s correlation to the panel mean for models trained using k-nearest neighbor (KNN) and RF, trained on cFPs or Mordred features, and the GNN model. Only the GNN model has a median correlation to the panel mean that is higher than that of the median panelist.

As a baseline comparison, we trained a cFP-based random forest (RF) model, the previous state-of-the-art (*6*), on the same dataset (Fig. 2B). This baseline model surpassed the median panelist for only 41% of molecules, showing that our GNN model’s performance increase comes not only from the volume and quality of the data, but importantly from the model architecture.

The GNN model shows human-level performance in aggregate, but how does it perform across perceptual and chemical classes? When we disaggregate performance by odor label, the model is within the distribution of human raters for all labels except musk and surpasses the median panelist for 32/55 labels (58%, Fig. 3A). This per-label view supports the view that the GNN model is superior to the previous state of the art model trained on the same data (paired 2-tailed t-test p=1.0e-7).

**Fig. 3.**
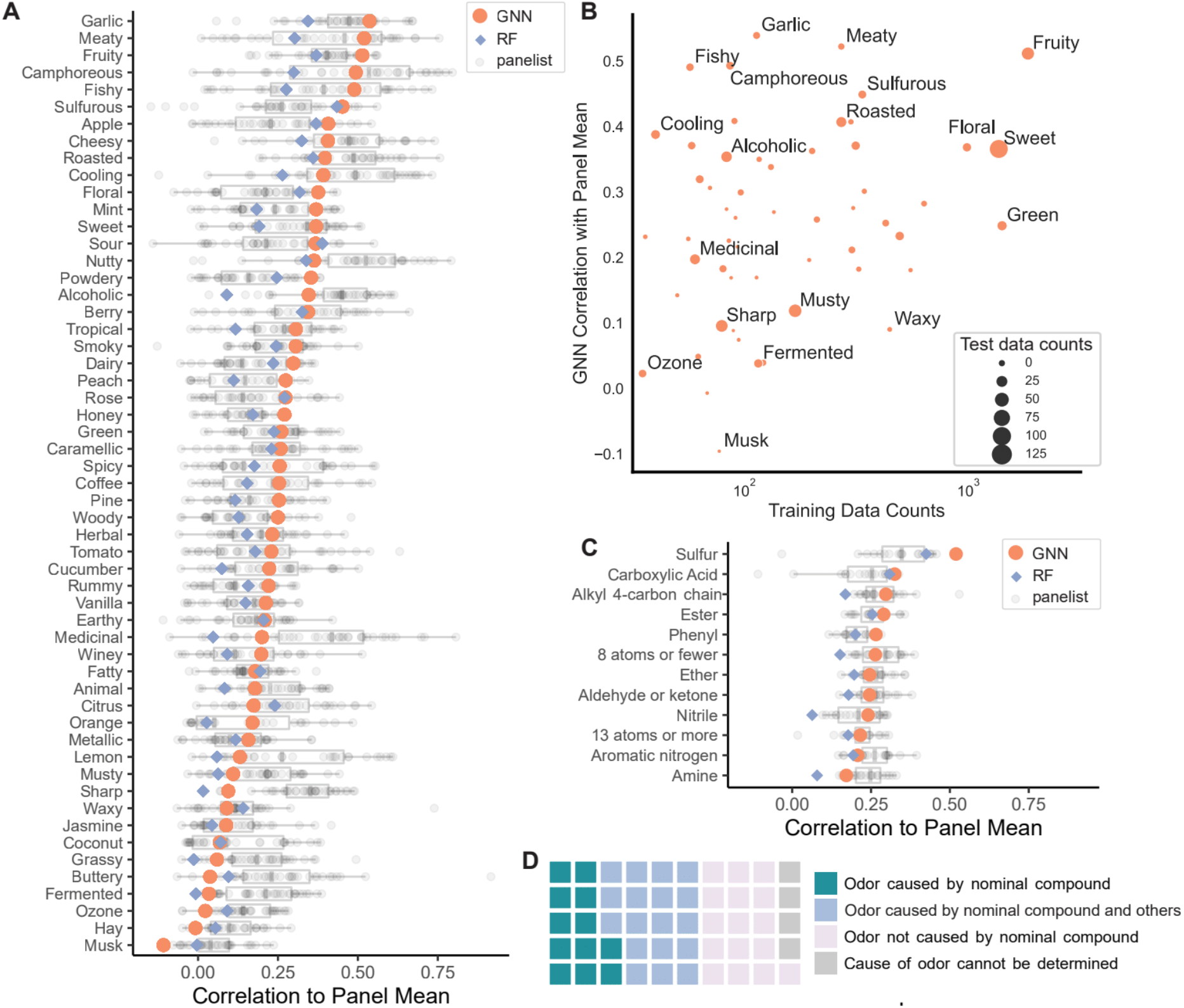
Model performance is robust across structural and perceptual classes. **(A)** Correlation of GNN (in orange) and RF (in purple) model predictions and panelist ratings (in gray) to the panel mean for each of the 55 odor labels. **(B)** GNN model correlation to panel mean for each of the 55 odor labels plotted against the number of molecules in the training data for which the label applies. Circle size is proportional to the number of test set molecules for which the label applies. Selected data points are annotated. **(C)** Mean correlation of GNN (in orange) and RF (in purple) model predictions and panelist ratings (in gray) to the panel mean for molecules belonging to 12 common chemical classes. **(D)** Categorization of gas chromatography-olfactometry quality control results for 50 validation set stimuli.

Predictive performance for a given label depends on the complexity of the structure-odor mapping for that label, so it is unsurprising that it performs best for labels like garlic and fishy that have clear structural determinants (sulfur-containing for garlic; amines for fishy), and worst for the label musk, which includes at least 5 distinct structural classes (macrocyclic, polycyclic, nitro, steroid-type, and straight-chain) (*21*, *22*). In contrast, a panelist’s performance for a given label depends on their familiarity with the label in the context of smell; consequently, we see strong panelist-panel agreement for labels describing common food smells like nutty, garlic, and cheesy and weak agreement for labels like musk and hay. Weak agreement for musk may also be due to genetic variability in perception, a well-documented phenomenon (*23*).

Model performance also depends on the number of training examples for a given label; with enough examples, models can learn even complex structure-percept relationships. In general, our model’s performance is high for labels with many training examples (e.g, fruity, sweet, floral) (Fig. 3B), but performance for labels with few training examples can be either high (e.g., fishy, camphoreous, cooling) or low (e.g, ozone, sharp, fermented). In other words, collecting more training data raises the floor for model performance. Likewise, model performance is bounded above by panel test-retest correlation (Fig. S13). When we disaggregate by chemical classes (e.g. esters, phenols, amines), both panelist and model performance is relatively uniform (Fig. 3C), with sulfur-containing molecules showing strongest performance from panelists and the model (R = 0.52).

Chemical materials are impure - a fact too often unaccounted for in olfactory research(24). To measure the contribution of impurities to the odor percept of our stimuli, we applied a gas chromatography-mass spectrometry (GC-MS and gas chromatography-olfactometry (GC-O) quality control (QC) procedure to 50 stimuli (Data S1). This QC procedure matches an odor percept to its causal molecule, allowing us to identify stimuli for which the primary odor character was not due to the nominal compound. Our QC led to diverse conclusions: the nominal compound caused the odor (12/50), the nominal compound and contaminants contribute to the odor (16/50), contaminants caused the odor (18/50), or the cause of the odor could not be determined (4/50) (Fig. 3D). In some cases, while we purchased a novel odorant, the dominant odorant was not novel; for example, the stimulus 4,5-dimethyl-1,3-thiazol-2-amine was described by the panel as buttery, sweet, and dairy, but this odor percept was attributed through QC to the contaminant diacetyl, a well-known buttery odorant. In another case, the purchased odorant, isobornyl methylacrylate, was described by the panel and the model as both piney and floral; however, through QC we determined that the nominal compound was floral only and that the piney aroma was due to the closely related compound, borneol, which was detected as a contaminant in the sample. Based on QC results, we removed 26 molecules known or suspected to have high degrees of odorous contamination (Data S1).

The prevalence of odorous contamination that we found demonstrates that it is not safe to assume that the odor percept of a purchased chemical is due to the nominal compound. The Flavor & Fragrance (F&F) industry is motivated to minimize odorous contaminants for commercially valued odorants, but there is no such incentive for non-F&F commodity chemicals. We stress the need for caution and diligence in expanding odor stimulus space.

Implications of each QC result on model performance are unique (Data S1). In some cases, the model performed well despite the presence of odorous contaminants. We estimate that, if these contaminants were removed from the rated samples, model performance improves in 6 of 50 scenarios, degrades in another 6 of 50 scenarios, remains neutral in 21 of 50 scenarios, and cannot be determined in 17 of 50 scenarios.

To test if the model is robust to discontinuities in structure-odor distances, we designed an additional challenge in which 41 new triplets (example in Fig. 4A) were constructed and validated by the panel (as in Fig. 1A). In each triplet, the anchor molecule is a known odorant, and is matched with one structurally similar and one structurally dissimilar novel odorant, and in which the more *structurally dissimilar* odorant is predicted to be the more *perceptually similar* of the two to the anchor. Our trained panelists were presented with the three odorants as a set and rated the perceptual distance between each of the molecules in the triplet (Fig. 4B). Confirming the model’s predictions -- counterintuitive under simpler structural models of odor -- our panelists generally rated the molecules as being more to an anchor molecule than the anchor’s structural neighbor (p < 2.2e-16, Fig. 4C). This significant result is further evidence that the POM overcomes discontinuities in the structure-odor relationship.

**Fig. 4.**
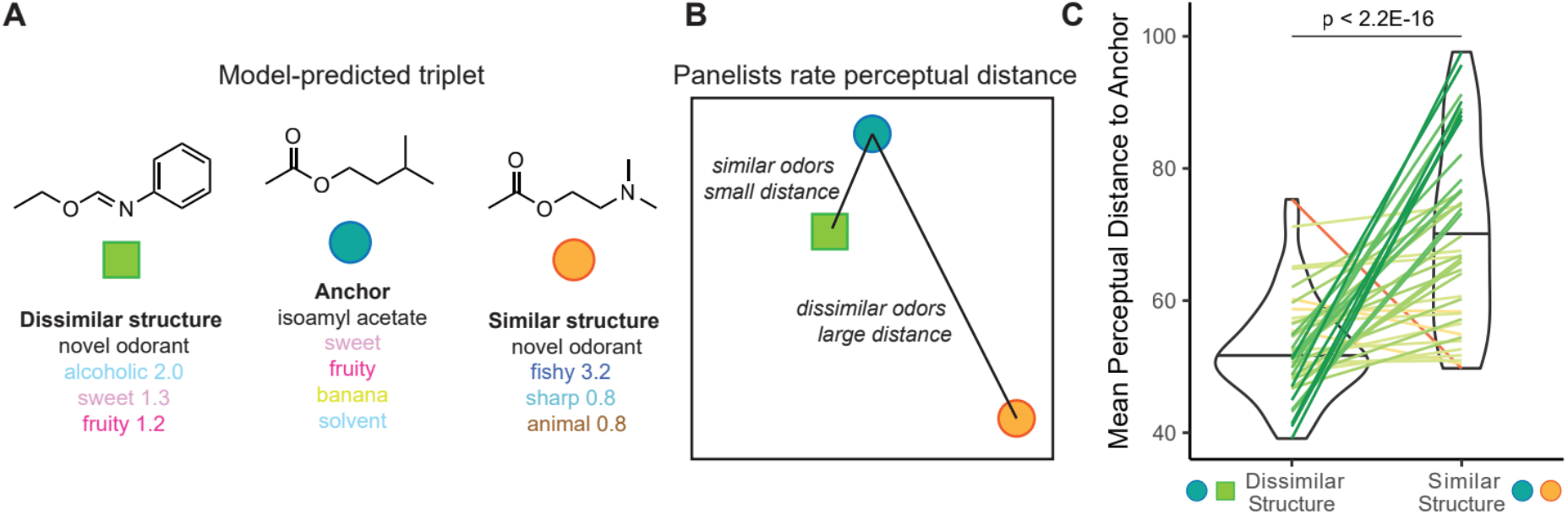
POM solves discontinuities in structure-odor mapping. **(A)** Example triplet of molecules identified by the GNN model in which the structurally similar pair is not the perceptually similar pair. We used the model to select 41 such triplets. **(B)** Diagram of the psychophysical task in which panelists rated perceptual distances between molecules in predicted triplets. **(C)** Mean perceptual distance rating for molecules that are structurally dissimilar (left) or structurally similar (right) to the same anchor molecule. Lines connect each pair of molecules compared to the same anchor molecule; line color corresponds to the relative difference in perceptual similarity. Perceptual similarity followed model predictions rather than structural similarity.

A reliable structure-odor map allows us to explore odor space at scale. We compiled a list of ~500,000 potential odorants whose empirical properties are currently unknown to science or industry; most have never been synthesized before. Because a molecule’s coordinates in the POM are directly computable from the model, we can plot these potential odorants in the POM (Fig. 5A), revealing a potential space of odorous molecules that is much larger than the much smaller space covered by current fragrance catalogs (~5,000 purchasable, characterized odorants). These molecules would take approximately 70 person-years of continuous smelling time to collect using our trained human panel.

**Fig. 5.**
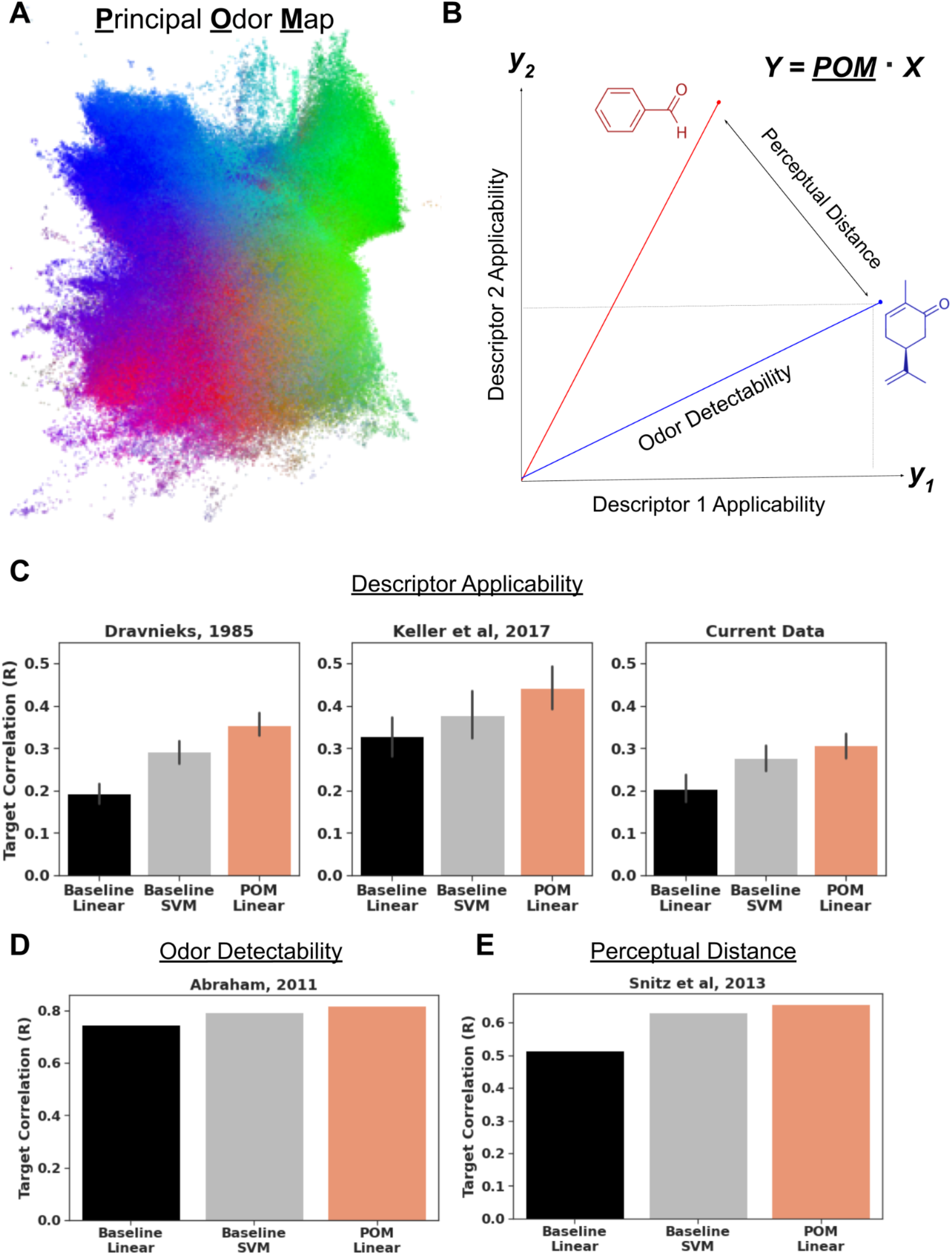
POM solves a fundamental set of olfactory prediction tasks. **(A)** 2D trimap embedding of 500,000 unique likely odorants previously uncharacterized. The position of each point (molecule) is determined by POM coordinates, and the RGB values of each point correspond to their coordinates in the first 3 dimensions of a non-negative matrix factorization of the predicted odor labels. **(B)** Intuitive geometric measures like vector length, vector distance, and vector projection correspond to the odor prediction tasks of odor detectability, similarity, and descriptor applicability. Equation shows that the projected space *Y* represents the dot product between POM and a task-specific projection matrix X. **(C)** A linear model atop POM outperforms a chemoinformatic SVM baseline at predicting odor applicability on two extant datasets, Dravnieks (*25*) and DREAM (*6*), as well as the current data. **(D)** A linear model atop POM outperforms a chemoinformatic SVM baseline at predicting odor detection threshold using data from Abraham et al, 2011 (*26*). **(E)** A linear model atop POM outperforms a chemoinformatic SVM baseline at predicting perceptual similarity on Snitz et al, 2013 (*4*).

We show that the POM has a meaningful interpretation by extracting intuitive, geometric measures and mapping them to several olfactory prediction tasks (Fig. 5B). The applicability of any set of odor descriptors corresponds to a projection of the POM coordinates onto axes corresponding to those descriptors; odor strength (detectability) corresponds to the magnitude of this projection (Fig. S12), and odor similarity corresponds to the distance between such projects for different molecules. We find that a simple linear model applied to POM and using these geometric interpretations has comparable or superior performance to a chemoinformatic support vector machine (SVM) model across multiple published datasets (Fig 5C, D, E), collectively representing some of the most thorough previous public efforts to characterize these features of odor.

## Discussion

There is no universally accepted method for quantifying and categorizing an odor percept. In other words, olfaction has been a sense without a map. Systems of odor classification have been proposed: first intuitive categorizations (*28*), then empirically-supported universal spaces (*29*, *30*), and later attempts to incorporate receptor mechanisms (*31*, *32*). However, these systems do not tie stimulus properties to perception, and none have reached broad acceptance. Here we propose and validate a novel, data-driven, high-dimensional map of human olfaction. We have shown that this map recapitulates the structure and relationships of odor perceptual categories evoked by single molecules, that it can be used to achieve prospective predictive accuracy in odor description that exceeds that of the typical individual human, and that it is broadly transferrable to arbitrary olfactory perceptual tasks using natural and interpretable transformations. This map represents for odor what the CIE color space represents for vision.

Nearly all published chemosensory models were fit to the data used in their construction. Even using cross-validation, the opportunity for over-fitting is high, because the data comes from a single distribution, task, or experimental source. Prospective validation on new data from a new source with no adjustments, as we performed, represents a much more stringent test of real-world utility. In this prospective context, we found that our model performs roughly on par with the median human panelist, beating a chemoinformatic baseline.

However, in a real-world setting, models can and should be updated as new data becomes available. This process is called ‘online learning’ (*27*), and is a central capability of many real-world ML systems. Fig 5C demonstrates that a linear model atop POM reaches an even higher level of performance when the POM is tuned to the new dataset.

The success of this model is not merely an advance in predictive modeling. It offers a simple, intuitive, contiguous, hierarchical, parseable map of molecular space in terms of odor, much as color spaces represents wavelengths of light in terms of colors and color components. It enables human-level performance not only for odor description but also generalizes to a gamut of other olfactory tasks. It offers the opportunity to reason, intuitively and computationally, about the relationships within and between molecular and odor spaces.

There are some practical considerations to keep in mind when using this map. First, the concentration of an odor influences odor character, but is not explicitly included in the map. So while it can predict detection thresholds, a property of the odorant molecule, it cannot predict suprathreshold intensity, a function of the odorant and its concentration. Many molecules have no odor, which we addressed by pre-screening with a separate, simpler model (*33*), and we diluted odorants to standardize intensity. Second, predictive performance is strong for organic molecules, the vast majority of odorants we encounter, but we could not extend the predictions into halides or molecules that include novel elements due to the lack of safety data for those molecules. Given uniformly strong performance across broad chemical classes tested in our prospective validation set (Fig. 3C), we expect high accuracy on novel chemicals within these chemical classes, but we would not expect high performance for molecules that have chemical motifs not represented in our training set. For instance, if our training dataset did not contain any molecules with carbon macrocycles, we would not expect the model to accurately predict the odor of an unseen macrocyclic musk (Fig. 3A). Third, many chemical stimuli have odorous contaminants (*24*), particularly those that have not been developed for use in fragrance applications. Neural networks are known to perform well, even with substantial noise in the training and test sets, which we see in the present work. Nonetheless, we recommend isolating the compound of interest from odorous contaminants, and/or characterizing the perceptual quality of contaminants. Finally, datasets in real-world settings are not static, but grow in size, and shift in distribution — models should be periodically retrained to incorporate new data. We showed that model performance tends to improve with increased training data (Fig. 3B) and data quality (Fig. S13), consistent with ML applications in other areas (*34*, *35*). Indeed, the most important future work -- work which will increase the accuracy and resolution of the map and any model that uses it -- will be scaling the volume and quality of training data.

Progress in neuroscience is often measured by the creation and discovery of new maps of the world supported by neural circuitry—maps of space in hippocampus, faces in the superior temporal sulcus, tonotopy in auditory cortex, and retinotopy and Gabor filters in V1 visual cortex, among others. Each is only possible because scientists first possessed a map of the external world, and then measured how responses in the brain varied with stimulus position on the map. We have had no such map for odor, but this study proposes and validates a novel data-driven map of human olfaction. We hope this map will be useful to researchers in chemistry, olfactory neuroscience, and psychophysics: first, as a drop-in replacement for chemoinformatic descriptors, and more broadly as a new tool for investigating the nature of olfactory sensation.

## Acknowledgements

The authors wish to acknowledge Zelda Mariet for experimental design, Yoni Halpern, Bob Datta, Steven Kearnes, Christina Zelano, Ari Morcos, Dan Bear, and Alex Koulakov for feedback on draft, contributions to GC-MS/O analysis from Dr. J.S. Elmore, and domain expertise from Christophe Laudemiel.

## Author contributions

Conceptualization: ABW; Methodology: BKL, EJM, RCG, JDM, BSL, JKP; Software: BKL, BSL, RCG, JNW; Validation: RCG, BKL; Formal analysis: BKL, EJM, RCG; Investigation: EJM, KAL, MA, BBN, TM, JKP, JDM, BSL, WWQ, JNW; Data curation: BKL, BSL, RCG, EJM, MA, KAL, JKP; Writing – original draft: EJM, BKL, ABW, JDM, RCG; Writing – review & editing: EJM, BKL, ABW, JDM, RCG, JKP, BSL, WWQ, JNW; Visualization: EJM, RCG, BKL, BSL; Supervision: ABW, JDM, EJM, JKP; Funding acquisition: ABW, JDM, EJM; Project administration: ABW, JDM

## Data and materials availability

Human psychophysics data, model predictions, and model embeddings for all 400 tested odorants are included, and will be deposited at the olfactory data repository pyrfume.org upon publication; odorant identities will be released pending legal review upon publication. Lightweight reproduction notebooks and scripts will be shared via https://github.com/google-research/google-research.

