## Supplemental Materials for "A Principal Odor Map Unifies Diverse Tasks in Human Olfactory Perception"

### Supplemental Methods

#### Training dataset

The GoodScents (http://www.thegoodscentscompany.com/) and Leffingwell PMP 2001 (https://zenodo.org/record/4085098#.YqoYk8jMlUE) datasets each contain odorant molecules and corresponding odor descriptors. Variations and misspellings of odor descriptors were merged, and any odor descriptor with <=30 occurrences in the dataset were discarded. The remaining list of odor descriptors is: [
'alcoholic', 'aldehydic', 'alliaceous', 'almond', 'amber', 'animal',
'anisic', 'apple', 'apricot', 'aromatic', 'balsamic', 'banana', 'beefy',
'bergamot', 'berry', 'bitter', 'black currant', 'brandy', 'burnt',
'buttery', 'cabbage', 'camphoreous', 'caramellic', 'cedar', 'celery',
'chamomile', 'cheesy', 'cherry', 'chocolate', 'cinnamon', 'citrus', 'clean',
'clove', 'cocoa', 'coconut', 'coffee', 'cognac', 'cooked', 'cooling',
'cortex', 'coumarinic', 'creamy', 'cucumber', 'dairy', 'dry', 'earthy',
'ethereal', 'fatty', 'fermented', 'fishy', 'floral', 'fresh', 'fruit skin',
'fruity', 'garlic', 'gassy', 'geranium', 'grape', 'grapefruit', 'grassy',
'green', 'hawthorn', 'hay', 'hazelnut', 'herbal', 'honey', 'hyacinth',
'jasmin', 'juicy', 'ketonic', 'lactonic', 'lavender', 'leafy', 'leathery',
'lemon', 'lily', 'malty', 'meaty', 'medicinal', 'melon', 'metallic',
'milky', 'mint', 'muguet', 'mushroom', 'musk', 'musty', 'natural', 'nutty',
'odorless', 'oily', 'onion', 'orange', 'orangeflower', 'orris', 'ozone',
'peach', 'pear', 'phenolic', 'pine', 'pineapple', 'plum', 'popcorn',
'potato', 'powdery', 'pungent', 'radish', 'raspberry', 'ripe', 'roasted',
'rose', 'rummy', 'sandalwood', 'savory', 'sharp', 'smoky', 'soapy',
'solvent', 'sour', 'spicy', 'strawberry', 'sulfurous', 'sweaty', 'sweet',
'tea', 'terpenic', 'tobacco', 'tomato', 'tropical', 'vanilla', 'vegetable',
'vetiver', 'violet', 'warm', 'waxy', 'weedy', 'winey', 'woody'
]
These datasets were merged and are subsequently referred to as “GS/LF”.

#### Model Training and Tuning

The network consists of several message-passing layers, followed by a radius 0 combination to fold atom and bond embeddings together, followed by a reduce-sum across atoms, followed by several fully connected layers and a final sigmoid function to make label predictions for each of the 138 curated descriptors described above.

All references to the GNN “embedding space” refer to the 256-dimensional activation of the final dense neural network layer. These embeddings are mapped to the final 138-dimensional prediction by one final dense layer of 138 neurons followed by a sigmoid function.

Hyperparameters of the neural network were optimized using 5-fold cross validation in our training set of ~4,000 molecules, using 500 trials of random search. Each model fit took less than 1 hour on a Tesla P100. We present results for the model with the highest mean AUROC on the cross-validation set. Since our multi-label problem had highly unbalanced labels, we used second-order iterative stratification to build our train/test/validation splits [47]. Iterative stratification is a procedure for stratified sampling that attempts to preserve many-order label ratios, prioritizing more unbalanced combinations. For second order, this means preserving ratios of pairs of labels in each split.

The objective function for training was a summed cross-entropy loss over all 138 descriptors, with each descriptor's contribution to the loss being weighted by a factor of log(1+ class_imbalance_ratio), such that rarer descriptors were given a higher weighting. l1 and l2-norm losses were also utilized.

For our random forest (RF) baseline methods, we tuned an exhaustive space of configurations of fingerprinting methods (bits, radius, counted/binary, RDKit/Morgan), and RF hyperparameters. The RDKit software(*36*) was used to calculate all features. We found a radius-4, 2048-bit Morgan fingerprint to perform most strongly in predicting odor labels.

Using an 80-20 stratified train/test split, we found that a trained GNN achieved an AUROC of 0.894 [CI 0.888 - 0.902] on the combined GS/LF dataset, whereas RF on Morgan fingerprints was the strongest baseline method with an AUROC of 0.850 [CI 0.838 - 0.860]. (*15*)

#### Model- and label-space comparisons

After applying a top-2 principal component reduction to GNN embedding, Morgan fingerprints, and perceptual label spaces, we found that GNN embedding inter-cluster distances correlate strongly ($r = 0.725$) with label inter-cluster distances, whereas Morgan fingerprints correlate weakly ($r = -0.119$). Inter-cluster distances are computed as the mean Euclidean distance of all |Cluster 1| * |Cluster 2| pairwise molecule distances. Fig S1A and S1B show all (138^2 - 138) pairwise comparisons of odor clusters.

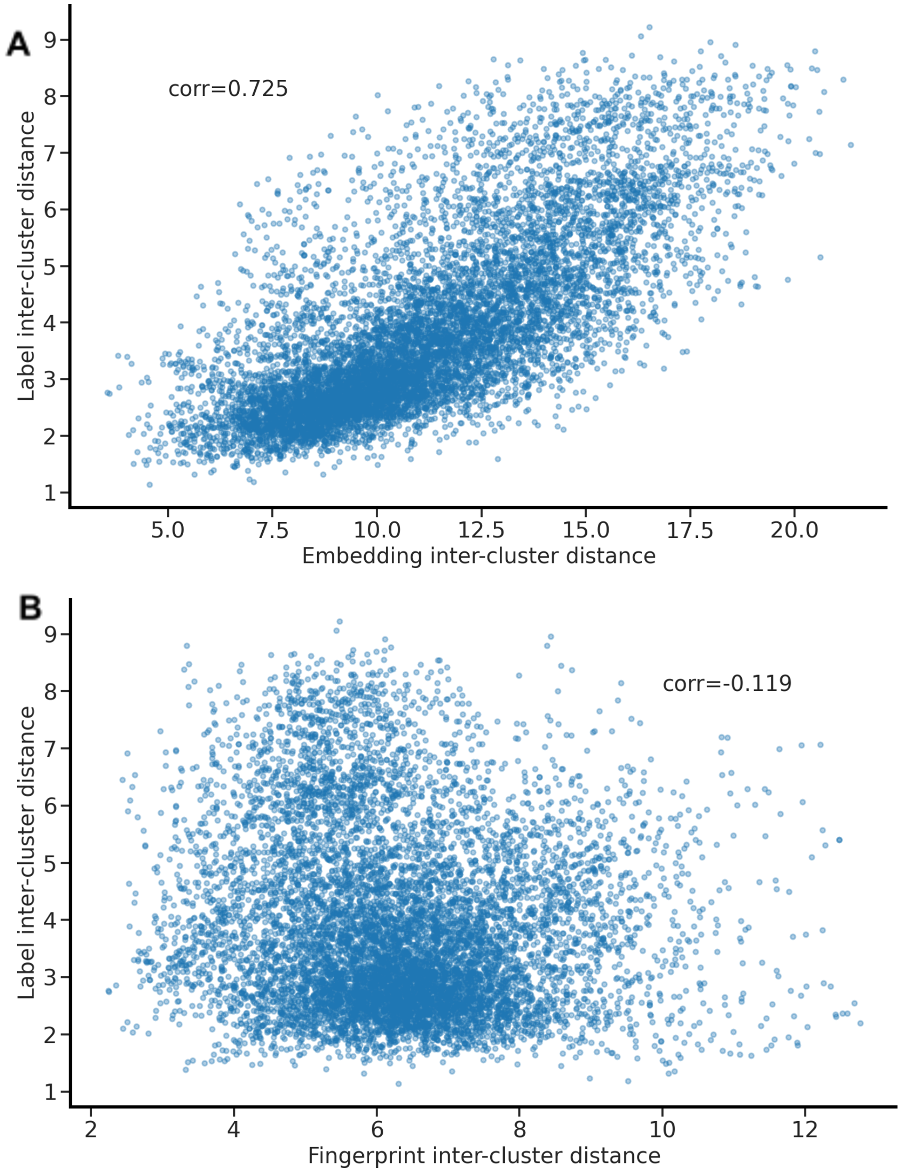


**Fig. S1.** Inter-cluster distance correlation. Each point represents a pair of odors (e.g. [fruity, sweet]. **(A)** Cluster distance, as measured by Euclidean distance of embeddings, correlates strongly with Jaccard overlap of clusters. **(B)** Cluster distance, as measured by Euclidean distance of fingerprints, does not correlate with Jaccard overlap of clusters.

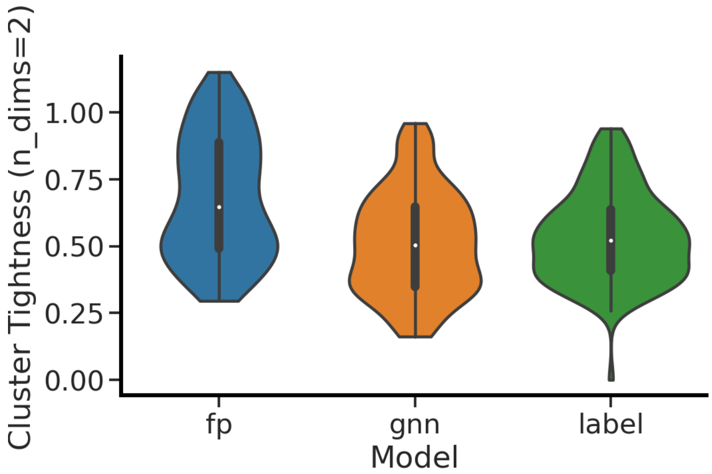


**Fig. S2.** Distribution of cluster tightness across 138 odor classes. Cluster tightness is defined as the ratio of {mean Euclidean distance of all in-group pairwise distances} to {mean Euclidean distance of all in/out-group pairwise distances}. 50th percentile cluster tightness for GNN embedding space was 0.513, similar to cluster tightness for label space (0.536), and tighter than fingerprint space (0.679).

#### Collecting a Prospective Validation Set

##### Aroma lexicon

We selected 55 of the 138 common labels from the GS/LF dataset to form our lexicon. We prioritized the selection of broad category terms (e.g., fruity, floral) to span the range of possible odor percepts, but also included specific terms from within an aroma category to measure model precision (e.g., fruity/citrus/lemon). Hierarchical clustering of GS/LF data supported our selections. Subjects used this lexicon exclusively to describe their odor quality perception of the odorants. The final list of odor descriptors used during human labeling is presented in Data S1.

**Table S1. Lexicon and associated aroma references.** The 55 descriptors in the lexicon were organized so that perceptually similar terms were close together in the list; terms are listed in the order they appeared on rater's ballots. Each term was paired with one or more aroma references to facilitate rater training.

| Lexicon Term | Aroma References |
|---|---|
| green | Le Nez Du Vin aroma standard - Vegetal |
| grassy | 2% cis-3-hexenol solution |
| cucumber | 1% nona-2,6-dienal solution |
| tomato | AromaMasters aroma standard - Tomato |
| hay | Animal bedding |
| herbal | Herbs de Provence; Dried dill |
| mint | Dried mint |
| woody | AromaMasters aroma standard - Cedar; Pine shavings |
| pine | 5% alpha-pinene solution |
| floral | AromaMasters aroma standard - Linden; AromaMasters aroma standard - Honeysuckle |
| jasmine | Jasmine essential oil |
| rose | Rose water |
| honey | Honey |
| fruity | Good & Gather flavor fusion fruit strips |
| citrus | Combined essential oils of orange, grapefruit, lime, and lemon |
| Lemon | Lemon essential oil |

| | |
|---|---|
| orange | Orange essential oil |
| tropical | Trident tropical twist gum |
| berry | Le Nez Du Vin aroma standard - Bilberry |
| peach | Le Nez Du Vin aroma standard - Peach |
| apple | Jolly Rancher - green apple |
| sour | White vinegar |
| fermented | GT's original kombucha |
| alcoholic | 200 proof ethanol |
| winey | Sutter Home red wine blend |
| rummy | Oakheart spiced rum |
| caramellic | Caramel flavor extract |
| vanilla | Vanilla extract |
| spicy | Blend of ground cinnamon, nutmeg, cloves, and allspice; Ground black pepper |
| coffee | Folgers medium roast ground coffee |
| smoky | 5% guaiacol solution |
| roasted | Le Nez Du Vin aroma standard - Toasted |
| meaty | Le Nez Du Vin aroma standard - Cooked beef |
| nutty | Roasted mixed nuts |
| fatty | 10% (E,E)-2,4-decadienal solution |
| coconut | AromaMasters aroma standard - Coconut |
| waxy | Crayola crayon |
| dairy | Carnation half & half pods |
| buttery | Butter extract |
| cheesy | Kernel Seasons white cheddar powder |
| sulfurous | Le Nez Du Vin aroma standard - Rotten egg |
| garlic | Garlic powder |
| earthy | Peat moss |
| musty | 0.1% 2,4,6-tribromoanisole solution |
| animal | AromaMasters aroma standard - Horse sweat |
| musk | Solution of galaxolide, ethylene brassylate, and tonalide |
| powdery | Johnson & Johnson baby powder |
| sweet | Charms cotton candy |
| cooling | Menthol crystals |
| sharp | 10% acetic acid solution |
| medicinal | Vicks VapoRub |
| camphoreous | 1% camphor solution |
| metallic | Pennies to be rubbed against skin |
| ozone | Adoxal |
| fishy | 0.5% trimethylamine solution |

**Panelist training and screening**

A pool of 26 prospective panelists between the ages of 18 and 55 and with a normal sense of smell were recruited from the Philadelphia area to participate in a 5-session series of training and screening exercises. The research protocol was approved by the University of Pennsylvania IRB, and all subjects gave informed consent prior to enrolling in the study. Subjects received odorant kits shortly before the start of the experiment and participated in sessions from home, facilitated by an experimenter over a Zoom video call (Zoom Meetings, https://zoom.us). The initial odorant kit (Fig. S3) contained 58 odor references (Table S1), 10 blinded odor references used for training quizzes, and 20 common odorants used in screening exercises (Data S2).

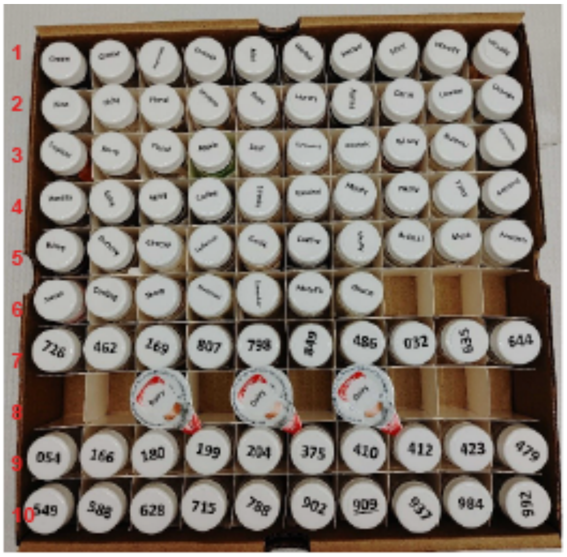

**Figure S3.** Initial odorant kit, containing 58 unique odor references corresponding to odor labels in the lexicon (rows 1-6, 8), 10 blinded odor references (row 7), and 20 blinded common odorants (rows 9-10).

In the first session, subjects were introduced to the study and trained to use the rate-all-that-apply (RATA) method to describe their perception of odorants (*37*); in the RATA method, subjects choose from a list terms that apply to the sample being evaluated, and then rate how strongly the chosen terms apply to the sample from 1 (low/slightly applicable) to 5 (high/very applicable). Subjects were given guidance on how to evaluate the odorants (e.g. take several short sniffs, hold the vial far from the nose to start and gradually bring it closer while sniffing, keep the cap tightly on each vial while not actively evaluating it) and taught to use a standard nasal rinse protocol (wet clean washcloth until just saturated, heat in microwave for 60s to produce steam, breathe in moist air above washcloth through nose for 30 seconds between evaluations, re-warm washcloth as needed; washcloths were provided with odor kits). Subjects then evaluated 20 common odorants using the RATA method as a pre-test.

In the second and third sessions, researchers trained subjects on the meaning of labels in the aroma lexicon. For each label, researchers described the olfactory meaning of the label, showed a related image, and prompted subjects to smell the associated odor reference(s). At the end of each session, subjects participated in a quiz in which they tried to identify blinded odor references and mixtures of references. Researchers then revealed the true identity of the blinded references and led a discussion about the results, prompting subjects to re-smell references and reinforce label meanings.

Following training, subjects evaluated the 20 common odorants in the fourth and again in the fifth session. Researchers reviewed subjects' quiz responses label selections for the 20 common odorants and calculated their test-retest correlation for post-training ratings. We invited 18 subjects who met our test-retest criterion (R>0.35) and made reasonable label selections for common odorants (e.g. mint selected to describe (-)-carvone) to join our panel (12 female, 6 male; 12 Caucasian, 4 African American or Black, 2 Asian; 3 Hispanic or Latino/a).

#### Virtual screening protocol for molecule selection

We began by filtering molecules listed in the eMolecules catalog -- which contains ~1 million commercially available molecules -- for atom composition (C/N/O/S/H only), price (<$1000 per 10 grams), purity (>95%), and availability (<4 weeks lead time). We developed a toxicity filter to conservatively remove potentially irritating or harmful compounds, (protocol developed by a certified toxicologist, approved by the University of Pennsylvania IRB), and removed likely odorless molecules according to water-soluble (cLogP < 0) and nonvolatile (boiling point > 300 C) criteria. We manually removed molecules that were likely to degrade or react under our experimental conditions. Finally, we compared predicted odor descriptors to the odor descriptors of all structurally similar reference molecules. All selected molecules satisfied one of two criteria:

1. Structurally similar to a molecule in the reference GS/LF dataset, yet with a negative prediction for that molecule's given descriptors. Prediction thresholds for descriptors were set at a threshold according to a geometric mean of training data frequency and test data empirical label frequency.
2. Structurally dissimilar to all molecules in the reference GS/LF dataset having a particular descriptor.

We selected and purchased 580 structurally distinct molecules from these structurally/perceptually divergent candidates (Fig. S4). Upon receipt of purchased molecules, we manually inspected for odorless molecules and diluted with propylene glycol to manually intensity-balanced each sample. 400 molecules were evaluated by human panelists, and the remainder of molecules were not tested further. Selection rationale for the 400 molecules is noted in Data S1.

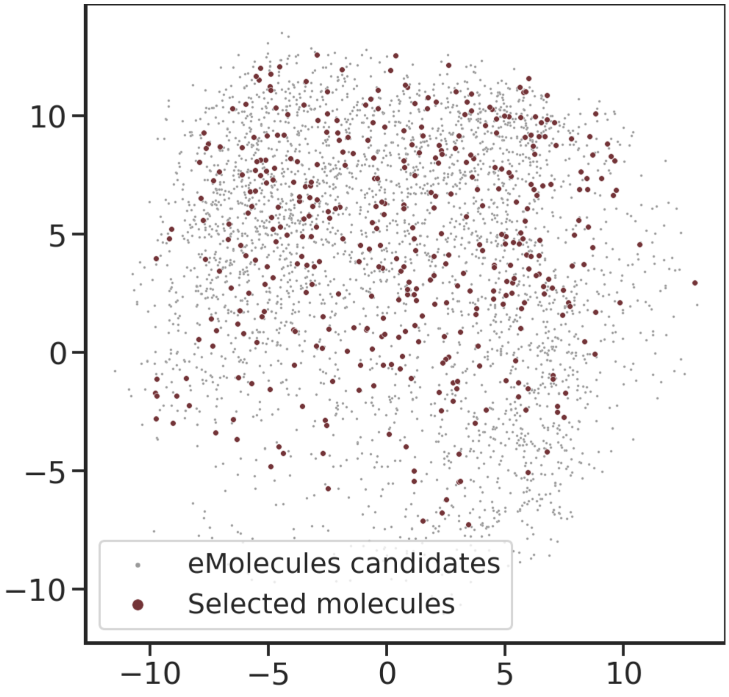


**Fig. S4**. Candidate and selected molecules from the eMolecules catalog displayed in chemical feature PCA space. Selected molecules span the space of potential candidate molecules from eMolecules.

**Odor evaluations**

Invited panelists were asked to rate the applicability of the 55 odor labels for each sample using the RATA method, as well as rate the intensity and pleasantness of each sample. Panelists received the odorants in sets of 50 and evaluated each twice over 4 sessions (25 evaluations/session). There was an enforced 30s break between each evaluation, and subjects followed the standard nasal rinse protocol described above during that break. In total, we characterized 400 novel odorants using this approach, and at least 15 of the initial 18 panelists participated in each phase of the study such that $n \geq 15$ for each odorant*replicate.

Against the backdrop of a global COVID-19 pandemic, we were wary of COVID-induced anosmia. Each session began with a training warm-up exercise to engage panelists in the rating task, reinforce label meanings, and enable researchers to verify that panelists had a normally functioning sense of smell. No subjects became anosmic during the course of the study.

Overall, we collected 400 molecules X 55 odor classes X 15 panelists X 2 replicates = 660,000 human sensory data points. The raw ratings are provided in Data S3, and summary statistics are described in Fig. S5. The distribution of non-zero descriptor ratings is shown in Fig. S5A, and the distribution of the number of descriptors applied to each molecule is shown in Fig. S5B. Each molecule is typically assigned between 1-6 descriptor ratings by each rater. Most descriptors are used at least once by every rater. Fig. S5C shows the percentage of molecules that are described by each of the 55 terms in the lexicon. Sweet was the most commonly applied descriptor.

Descriptor ratings show a clear correlation structure (Fig. S6). For example, fruity descriptors are more likely to co-occur with each other and less likely to co-occur with other descriptors including meaty, sulfurous, and roasted. Descriptor ratings are also related to odorant chemical class (Fig. S7). For example, molecules containing a sulfur atom are more likely to be described as meaty, molecules containing an amine group are more likely to be described as fishy, and molecules containing a carboxylic acid group are more likely to be described as sour.

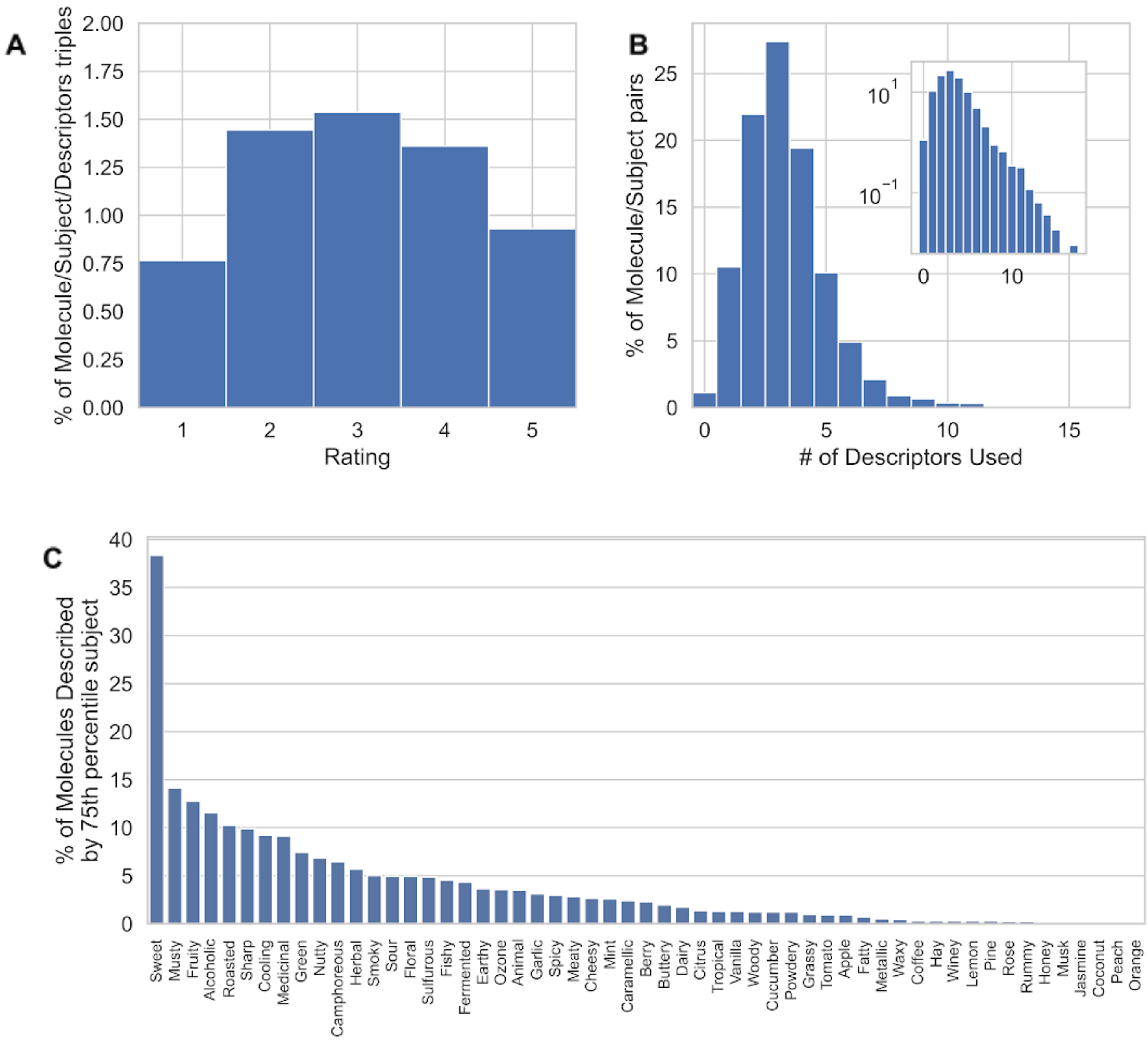


**Fig. S5. Human psychophysics prospective validation set summary statistics. (A)** Distribution of non-zero descriptor ratings, **(B)** distribution of the number of descriptors applied to each molecule, and **(C)** percent of molecules described by each of the 55 odor descriptors according to the 75th percentile panelist's ratings.

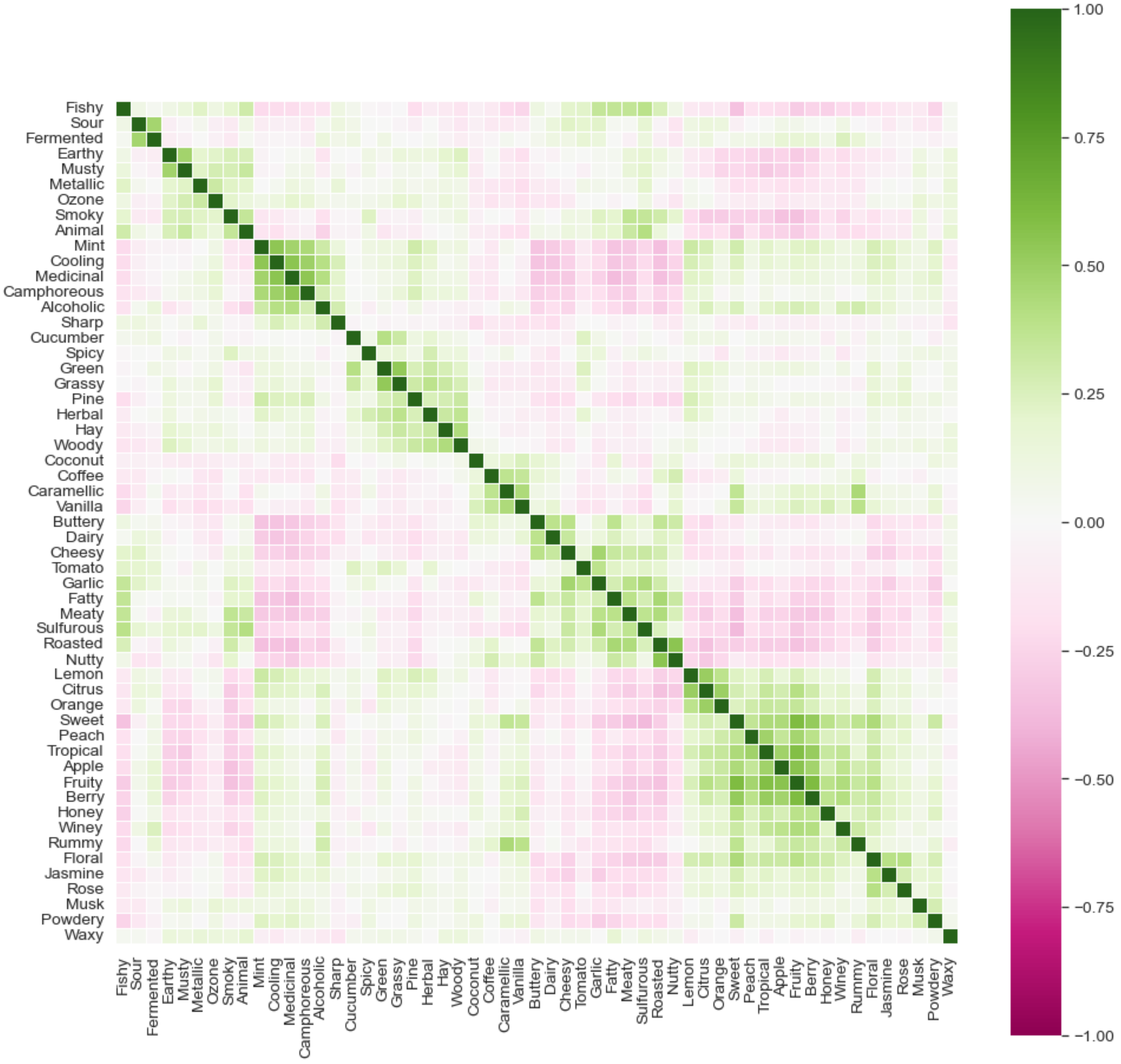


**Fig. S6.** Correlation matrix of panelist ratings for the 55 odor lexicon descriptors. Descriptors with strong positive correlations are dark green; descriptors with strong negative correlations are in dark pink. Odor descriptors show a clear correlation structure.

| | Sulfur | Amine | Phenyl | Nitrile | Ester | Carbonyl | Carboxylic Acid | Alkyl | <=8 atoms (<25%ile) | >=13 atoms (>75%ile) |
|---|---|---|---|---|---|---|---|---|---|---|
| roasted_cluster | 0.184 | 0.187 | 0.054 | 0.064 | 0.080 | 0.064 | 0.061 | 0.045 | 0.212 | 0.113 |
| meaty_cluster | 0.244 | 0.177 | 0.083 | 0.039 | 0.153 | 0.088 | 0.068 | 0.069 | 0.171 | 0.134 |
| fishy_cluster | 0.020 | 0.301 | 0.022 | 0.033 | 0.037 | 0.068 | 0.000 | 0.162 | 0.065 | 0.059 |
| primeval_cluster | 0.071 | 0.203 | 0.280 | 0.031 | 0.056 | 0.088 | 0.054 | 0.130 | 0.105 | 0.082 |
| gourmand_cluster | 0.000 | 0.018 | 0.050 | 0.209 | 0.043 | 0.078 | 0.054 | 0.000 | 0.072 | 0.026 |
| herbal_cluster | 0.010 | 0.062 | 0.224 | 0.088 | 0.048 | 0.067 | 0.021 | 0.113 | 0.072 | 0.112 |
| fruity_cluster | 0.006 | 0.107 | 0.234 | 0.113 | 0.204 | 0.178 | 0.054 | 0.088 | 0.108 | 0.181 |
| floral_cluster | 0.020 | 0.017 | 0.161 | 0.107 | 0.107 | 0.039 | 0.021 | 0.082 | 0.010 | 0.111 |
| sour_cluster | 0.037 | 0.059 | 0.037 | 0.015 | 0.061 | 0.064 | 0.262 | 0.064 | 0.096 | 0.068 |
| cooling_cluster | 0.008 | 0.148 | 0.148 | 0.055 | 0.114 | 0.160 | 0.043 | 0.118 | 0.154 | 0.065 |

**Fig S7.** Correlation of structural and perceptual categories. Each of the 400 molecules in the human validation set is non-exclusively classified according to structural and perceptual categories, and each table entry represents the Jaccard overlap (intersection over union) of molecule sets.

**Rater performance**

Aggregating all molecules and descriptors, our panel exhibited a test-retest correlation of 0.8. The panel test-retest correlation was high for most descriptors (Fig. S8). Compared to a prior large-scale human psychophysical study (*6*), our collected dataset has more descriptors and higher panel mean test-retest reliability (Fig. S9 and Fig. S10), even with fewer panelists.

Molecules with lower rated intensity were found to have the weakest panel test-retest correlations, indicating that the panel was not able to get a consistent evaluation (Fig. S11). We therefore excluded any molecules with intensity <3 (on a 0-10 scale) from the study. Of the 400 molecules evaluated by the panel, 42 were dropped from the validation set due to low odor intensity.

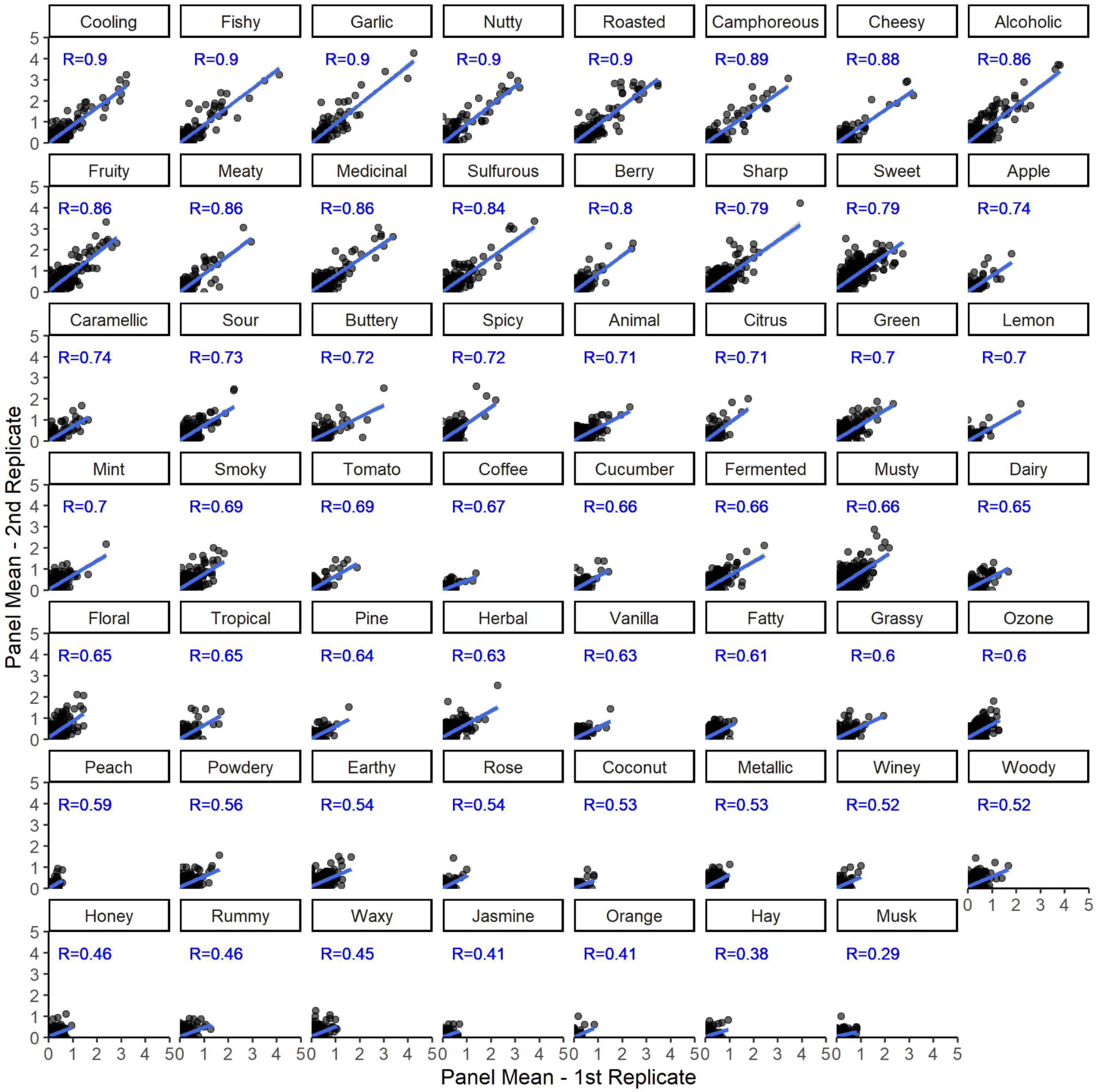


**Fig. S8.** Panel mean (n≥15 subjects) test-retest correlation (R) for the 55 descriptors in the lexicon applied to the 400 novel odorants in the prospective validation set. Descriptors are ordered by descending correlation.

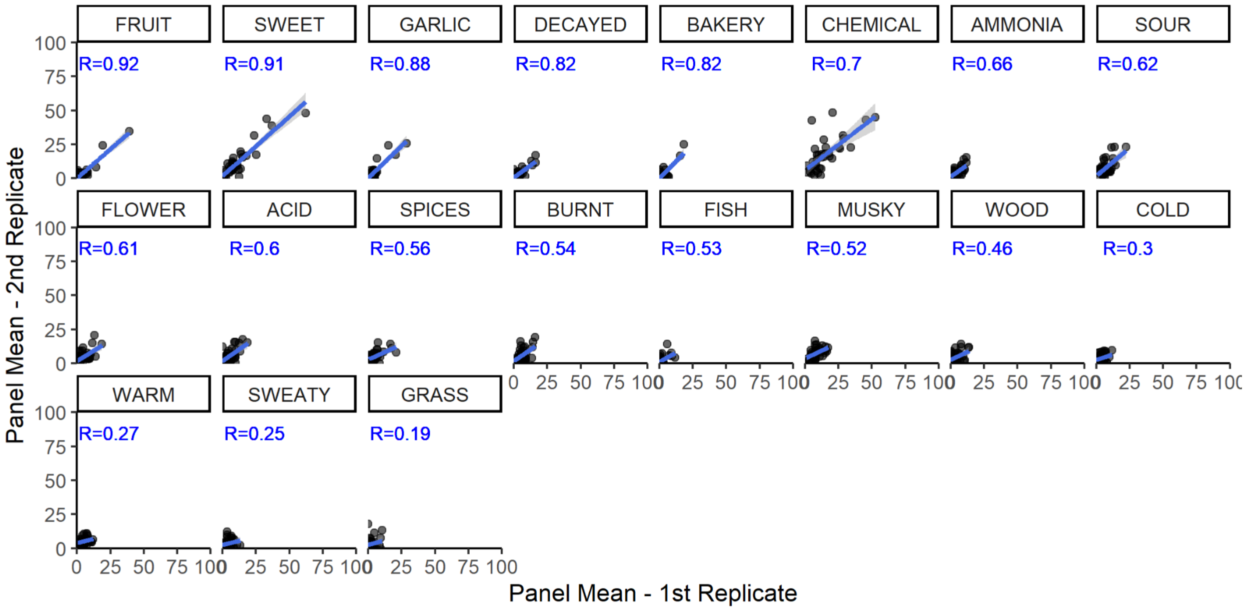


**Fig. S9.** Panel mean (n=49) test-retest correlation (R) for the 19 descriptors in the DREAM olfaction challenge dataset (6). Descriptors are ordered by descending correlation.

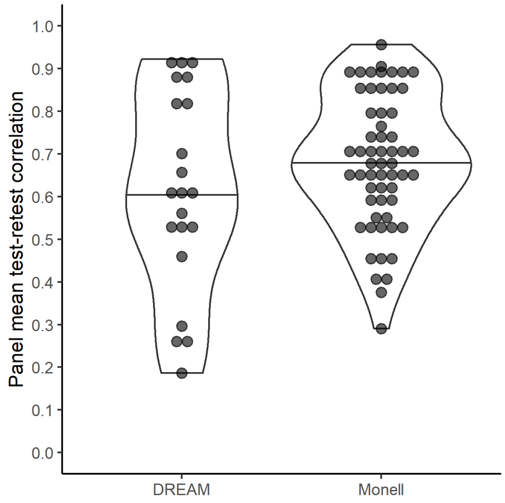


**Fig. S10**. Panel mean test-retest correlation for the 19 descriptors in the DREAM olfaction challenge (left) (6) and for the 55 descriptors in the present study (right). Each dot represents one odor descriptor.

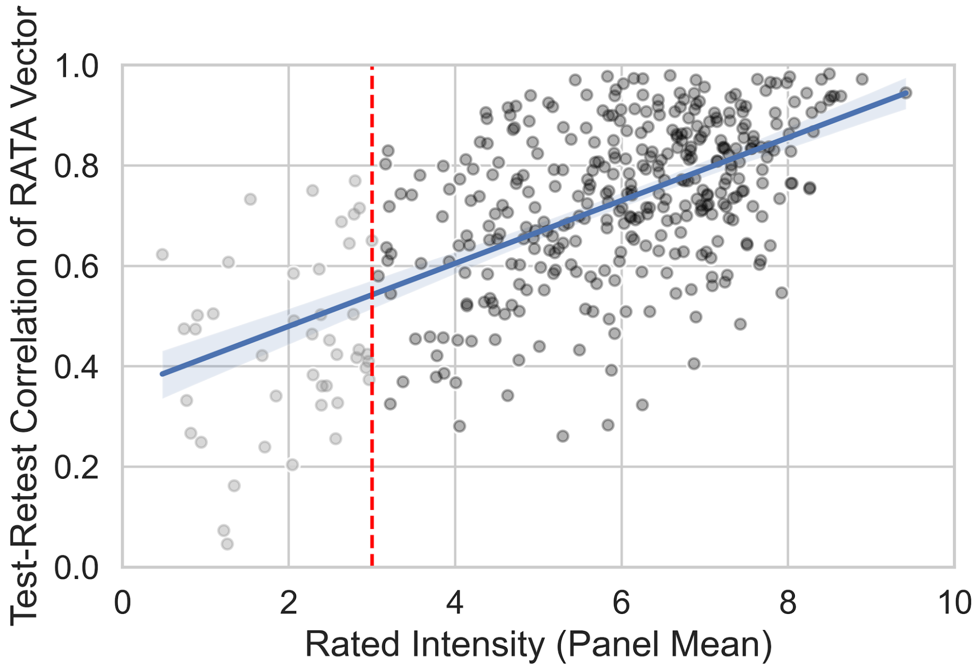


**Fig S11.** Test-retest correlation for 400 novel odorants as a function of panel mean intensity rating for that odorant. Molecules with lower rated intensity have weaker test-retest correlation of the panel mean.

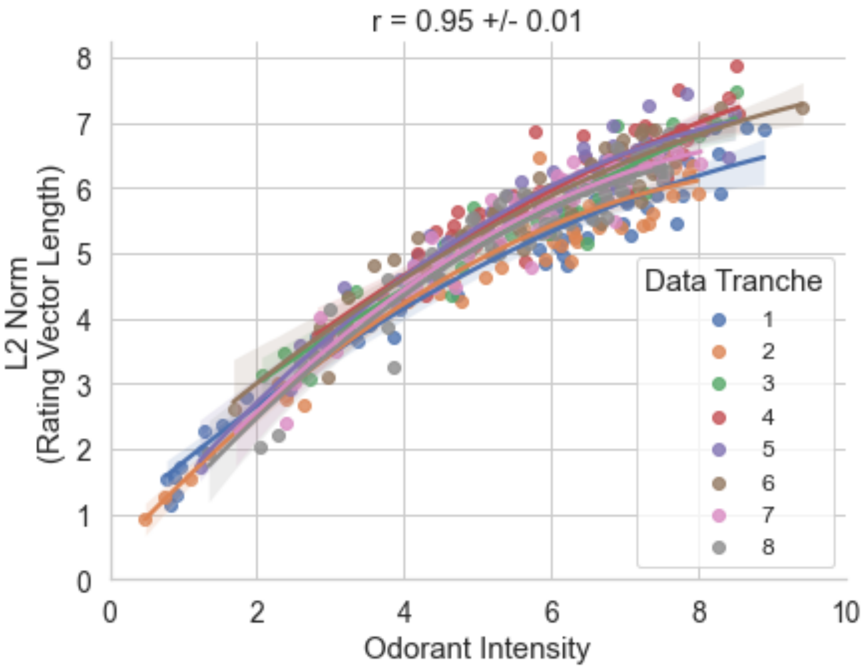

**Fig. S12.** L2 norm length of the mean RATA rating vector as a function of panel mean rated odorant intensity. Human psychophysical ratings were gathered in 8 data collection waves; differently colored dots and fits come from different tranches of data collection. Panelists use more descriptors and give higher RATA ratings for higher-intensity stimuli.

#### Evaluating Model Performance on Prospective Validation Set

We chose cosine similarity of a 55-dimensional vector as a metric that would emphasize overall accuracy of the predicted odor profile, rather than a “hit rate” on individual descriptors. This metric encountered some initial difficulties, as we discovered that the model and panelists were not directly comparable. The model makes an independent prediction for all 138 odor classes, resulting in a dense vector, whereas panelists typically rated only the top 3.2 ± 1.7 labels per odorant, resulting in sparse vectors. When shuffled, the model’s predictions had a positive nonzero score, indicating a systematic scoring bias in the model’s favor. We found that subtracting each individual model or panelists’ mean rating across all molecules from the respective predictions had the effect of zeroing out the shuffled baseline’s cosine similarity scores. We used this centered prediction or rating as the input to all of our calculations, as it would be a fairer comparison between model and panelist. Mathematically, this is similar to a Pearson correlation calculation, as the ratings are centered, but different due to not rescaling, as this would have destroyed useful information.

One disadvantage of the cosine metric is that it treats all 55 dimensions equally, yet not all mistakes are equally wrong. Descriptors have hierarchical relationships, and as such a “partial credit filter” -- which spreads observed single descriptor ratings across multiple descriptors -- can be learned and indeed can substantially improve performance (data not shown), but complicates the presentation of the results as it goes beyond simple arithmetic operations on raw data.

Model predictive performance is higher when human validation data has greater inter- and intra-subject agreement (Fig. S13). Increasing rater test-retest reliability and agreement is a necessary precursor to increasing measured model accuracy.

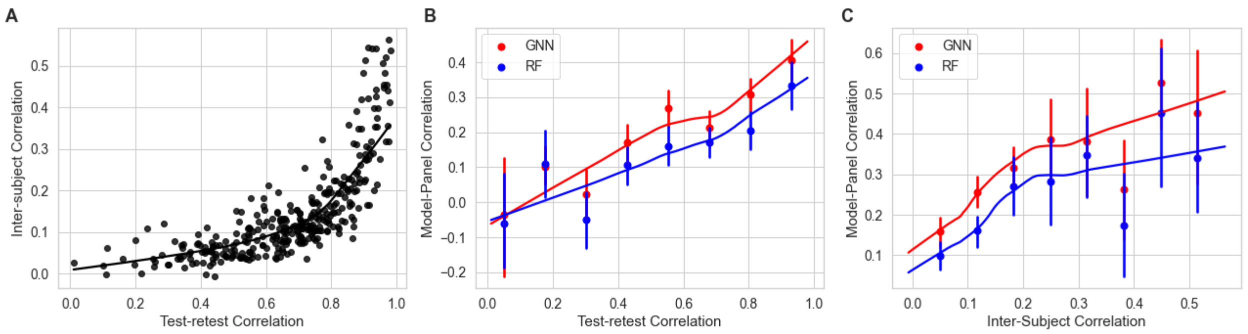


**Fig. S13.** Relationship between model predictive performance and inter- and intra-subject agreement. **(A)** Inter-subject correlation as a function of intra-subject correlation (test-retest correlation). **(B)** RF and GNN model-panel correlation as a function of binned test-retest correlation. **(C)** RF and GNN model-panel correlation as a function of binned inter-subject correlation. Model performance is capped by panelists' rating consistency.

#### Accounting for odorous contaminants

To account for the potential presence of odorous contaminants in the 400 commercial compounds purchased for the human validation study, we developed a gas chromatography-mass spectrometry/olfactometry quality control (QC) procedure. Fifty of the 400 molecules were selected for QC and shipped to the University of Reading for GC-MS/O analysis. By comparing retention indices of recorded odor percepts measured via GC-O to compound identities determined via GC-MS, we were able to identify cases in which contaminants influenced the odor of the material. We classified the molecules into one of 4 verdict categories: 1) Clean - no odorous contaminants found, 2) Mixed - odorous contaminant found but both nominal compound and contaminant contribute to odor, 3) Contaminated - odorous contaminant found, contaminant is the dominant contribution to odor, 4) Inconclusive - the causal odorant was not identified in GC-O, nor was there any detected odor at the expected elution time. This can happen due to thermal or oxidative degradation of the molecule under GC-O conditions, synergistic odorant combinations, or other experimental difficulties. GC-O experimenter notes and classification verdicts for the 50 QC-set molecules are included in Data S1.

In both QC-set cases where a non-sulfur containing molecule was rated sulfurous by the panel, GC-O showed that a sulfur-containing contaminant was the culprit. Additionally, in most QC-set cases where a non-dimethylamino-containing molecule was rated strongly fishy by the panel, GC-O showed that a dimethylamine contaminant was present. On this basis, molecules with an unexpectedly monotonic fishy/garlic/sulfurous profile were excluded from our analysis, including some molecules that had not been confirmed to be contaminated by GC-O analysis. Next, based on anecdotal reports from fragrance chemists that Michael acceptors are aggressive nucleophile scavengers, we excluded Michael acceptors that were reported as garlicky (phosphorous or sulfur impurity), sulfurous (sulfur impurity) or fishy (nitrogen impurity). Acetylene derivatives are also often garlicky due to phosphine ($PH_3$) impurities and we excluded a few molecules fitting this profile. In total, 26 molecules were dropped from the validation set due to confirmed or potential contamination. The rationale for these exclusions are included in table S1. The decision to exclude these molecules had no significant impact on model performance.

#### GC-O and GC-MS procedures

**Extraction of the compound onto the fiber:** The 50 compounds destined for gas chromatography-olfactometry (GC-O) and gas chromatography-mass spectrometry (GC-MS) were supplied either diluted in polyethylene glycol or neat, and absorbed onto Viscopearls (4 mm diameter, Rengo Co., Ltd., Osaka, Japan). For GC-O, approximately 10 balls (or fewer if the compound was very strong, more if it was very weak) were placed in a 20 mL SPME vial and equilibrated in a water bath prior to extraction onto a preconditioned triple phase solid phase microextraction (SPME) fiber (50/30 µm divinylbenzene/carboxen on polydimethylsiloxane (Supelco, Poole, UK). Generally, the samples were incubated at 45 or 55 °C depending on their volatility for 10 min, and extracted for a further 10 min (details in Data S1).

**Gas Chromatography-Olfactometry (GC-O):** After extraction, the SPME device was inserted into the injection port of an HP7890 GC from Agilent Technologies (Santa Clara, CA, USA) coupled to a Series II ODO 2 GC-O system (SGE, Ringwood, Victoria, Australia). The SPME fibre was desorbed in a split/splitless injection port held at 280 °C. The column employed was an Agilent HP-5 MSUi capillary (30 m, 0.25 mm i.d., 1.0 µm df) non-polar column. The temperature gradients was as follows: 40 °C initial temperature with a rise of 8 °C/min up to 200 °C and 15 °C/min from 200 °C to 300 °C and the final temperature held for a further 10 min. Helium was used as carrier gas (2 mL/min). At the end of the column, the flow was split 1:1 between a flame ionisation detector (kept at 250 °C) and a sniffing port using 2 untreated silica-fused capillaries of the same dimensions (1 m, 0.32 mm i.d.).

**Odor assessment**: Odor assessment was carried out by two flavour experts with 20 years' experience in using the GC-O, who had been familiarised with the standard lexicon. One expert assessed 46, compounds while the second assessed the remaining 4, and confirmed the assessment of a further 24 where clarification was required. Each assessor waited until the solvent had eluted (~5 mins) and sniffed the compounds eluting from the column until 20 min (equivalent to an LRI of 1700). They noted the time, intensity and descriptors for each compound that was detected. Linear retention indices were calculated by comparison with the retention times of C6-C25 n-alkane series analysed on the same day using the same conditions as for sample analyses. Where the LRI matched that of the target compound as determined by GC-MS, this was deemed to be the target compound and any other odors detected were contaminants.

**Gas chromatography-mass spectrometry (GC-MS):** For identification of the target compound by GC-MS, 2 balls (or more of it was very weak) were placed in a 20 mL SPME vial and equilibrated in a water bath at 30 °C for 10 min prior to a 30 s extraction onto the same SPME fibre type as used for GC-O.  For six less volatile samples (133 136 316 728 917), 10 balls were used, the incubation time was increased to 20 min at 55 °C, and extraction time increased to 20 min. A 7890A Gas Chromatograph coupled to a 5975C series GC/MSD from Agilent was used, equipped with the same column as described above. The oven started at 40 °C and increased to 300 °C at a rate of 8 °C/min. Helium was the carrier gas at a flow rate of 0.9 mL/min. Mass spectra were recorded in electron impact mode at an ionization voltage of 70 eV and source temperature of 220 °C. A scan range of m/z 25-450 with a scan time of 0.69 s was employed and the data were controlled and stored by the ChemStation software (Agilent, Santa Clara, CA). Linear retention indices were calculated by comparison with the retention times of C6-C25 n-alkane series analysed on the same day using the same conditions as for sample analyses. Compounds and contaminants were identified by comparison of their mass spectrum with those in the NIST 2020 library and, where available, the LRI was compared to that reported in the online NIST chemistry webbook or PubChem.

#### Historical explanations of Odor

Historical structure-odor relation models came in the form of empirical rules that are phrased as boolean logic expressions on the presence, absence, or proximity of molecular fragments.For example, Boelens' Rose rule is phrased as "the presence of a 7-9 carbon moiety with a hydroxy or oxy carbonyl or ether group attached to the moiety."(*38*) We also note that the original expression of these rules are often underspecified, meaning that these plain-language rules cannot be converted directly into e.g. Python code. We show two examples below, including Boelens' 1973 rose rule and Stoll's 1936 musk rule (Fig. S14)(*38–40*).

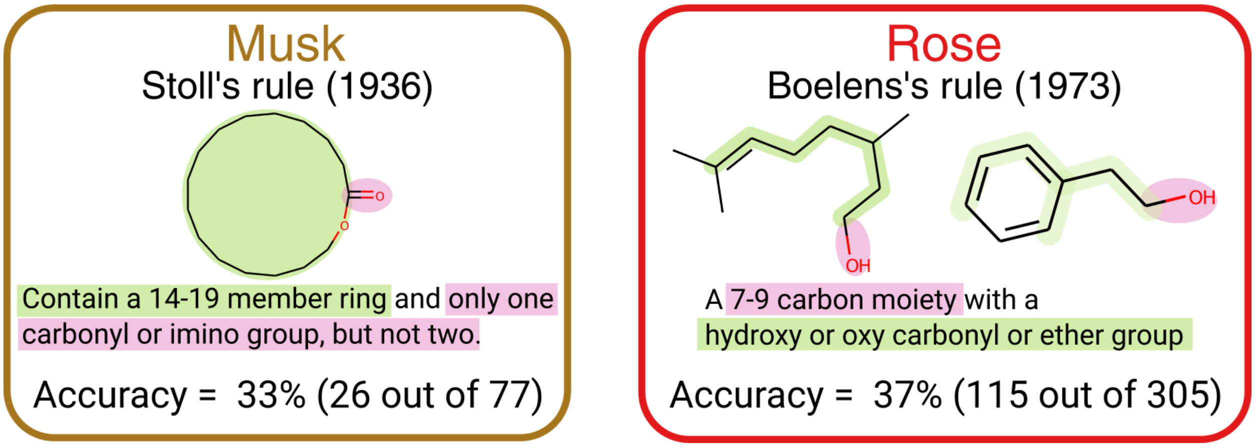


**Fig. S14:** Example of two historical odor rules with their recall as measured on the Goodscents, Leffingwell datasets.

### Supplemental Data

Molecules are indexed by a unique identifier (RedJade Code) allowing for reproduction of most results shown here. Information required to identify chemical structures will be provided upon publication.

**Data S1.** Metadata for 400 molecules comprising the prospective validation dataset. Columns in the dataset are defined as follows:

| | |
|---|---|
| RedJade Code | Internal anonymizing tracking number for panelist responses [PRIMARY KEY] |
| Odor Key | Tracking number from chemical inventory system, ignore. |
| Kit | Batch number of the molecule. Molecules were tested in 8 waves of 50 molecules. |
| Solvent | Diluting solvent, if needed for safety or for intensity balancing |
| Final [] | Concentration of molecule (w/w) in final sample |
| GC-O no of balls used | Number of viscopearls with absorbed odorant used for headspace extraction prior to GC-O analysis |
| GC-O incubation and extraction temperature | GC-O methodological details |
| GC-O incubation time | |
| GCO extraction time | |
| LRI GCMS | Linear Retention Indices for nominal compound via GCMS, if detected |
| LRI GC-O FID | Linear Retention Indices for nominal compound via GC FID, if detected |
| LRI GC-O aroma | Linear Retention Indices for odor of nominal compound via GC-O, if detected |
| GCO raw commentary | Raw notes from GC-O analyst, if molecule was tested with GC-O |
| GCO result | Verdict from GC-O analysis |
| GCO contaminant, if identified | Canonical SMILES of the causal contaminant, if |

| | |
|---|---|
| | one was successfully identified |
| Impact on GNN performance | Whether the GCO result had a good, bad, neutral, or unknown effect on GNN's prediction performance. |
| Disqualification reason | Reason for disqualification. If blank, molecule was retained for analysis |
| Selection reason | Original selection criteria. Molecules were predicted by the GNN or Random Forest model to have an odor prediction above some threshold despite structural dissimilarity to known instances of that odor class, or to have an odor prediction below some threshold, despite structural similarity to known instances of that odor class. |

**Data S2.** Panelist evaluations of 20 common odorants. Prospective panelists gave RATA ratings using the 55-word lexicon for 20 common odorants; panelists with a raw test-retest correlation greater than 0.35 were invited to join the panel.

**Data S3.** Panelist evaluations of 400 novel odorants. Between 15 and 18 panelists rated intensity and pleasantness and gave RATA ratings using the 55-word lexicon for each molecule.

**Data S4.** Odor attribute predictions on 400 molecules by a random forest model trained on GS/LF datasets.

**Data S5.** Odor attribute predictions on 400 molecules by a graph neural network model trained on GS/LF datasets. Final layer.

**Data S6.** Graph neural network embeddings on 400 molecules. Penultimate layer.

**Data S7.** Correspondence table between internal odorant identifiers and chemical structures.
